# Protein Vaccine Induces a Durable, More Broadly Neutralizing Antibody Response in Macaques than Natural Infection with SARS-CoV-2 P.1

**DOI:** 10.1101/2021.09.24.461759

**Authors:** Albert To, Teri Ann S. Wong, Michael M. Lieberman, Karen Thompson, Laurent Pessaint, Jack Greenhouse, Nisrine Daham, Anthony Cook, Brandon Narvaez, Zack Flinchbaugh, Alex Van Ry, Jake Yalley-Ogunro, Hanne Andersen Elyard, Chih-Yun Lai, Oreola Donini, Axel T. Lehrer

## Abstract

FDA-approved and Emergency Use Authorized (EUA) vaccines using new mRNA and viral-vector technology are highly effective in preventing moderate to severe disease, however, information on their long-term efficacy and protective breadth against SARS-CoV-2 Variants of Concern (VOCs) is currently scarce. Here we describe the durability and broad-spectrum VOC immunity of a prefusion-stabilized spike (S) protein adjuvanted with liquid or lyophilized CoVaccine HT™ in cynomolgus macaques. This recombinant subunit vaccine is highly immunogenic and induces robust spike-specific and broadly neutralizing antibody responses effective against circulating VOCs (B.1.351 [Beta], P.1 [Gamma], B.1.617 [Delta]) for at least 3 months after the final boost. Protective efficacy and post-exposure immunity were evaluated using a heterologous P.1 challenge nearly 3 months after the last immunization. Our results indicate that while immunization with both high and low S doses shorten and reduce viral loads in the upper and lower respiratory tract, a higher antigen dose is required to provide durable protection against disease as vaccine immunity wanes. Histologically, P.1 infection causes similar COVID-19-like lung pathology as seen with early pandemic isolates. Post-challenge IgG concentrations were restored to peak immunity levels and vaccine-matched and cross-variant neutralizing antibodies were significantly elevated in immunized macaques indicating an efficient anamnestic response. Only low levels of P.1-specific neutralizing antibodies with limited breadth were observed in control (non-vaccinated but challenged) macaques suggesting that natural infection may not prevent reinfection by other VOCs. Overall, these results demonstrate that a properly dosed and adjuvanted recombinant subunit vaccine can provide long-lasting and protective immunity against circulating VOCs.

**One Sentence Summary:** A recombinant subunit protein formulated with CoVaccine HT™ adjuvant induces superior immunity than natural infection and reduces viral load while protecting cynomolgus macaques from COVID-19-like disease caused by late SARS-CoV-2 P.1 (Gamma) challenge.

## INTRODUCTION

COVID-19 is a respiratory disease caused by the Severe Acute Respiratory Syndrome Coronavirus 2 (SARS-CoV-2) (*1, 2*) and is commonly characterized as an asymptomatic infection or a self-limiting, febrile illness co-presenting with cough, shortness of breath, and fatigue (*3, 4*). Severe complications can result in approximately 6-10% of infected patients (*5*) developing pneumonia, acute respiratory distress syndrome, multi-organ dysfunction and arterial thromboembolic events that can result in hospitalization and/or death (*6-9*). Transmission occurs through respiratory droplets during close contact with asymptomatic or presymptomatic infected individuals (*10-12*). The synergistic relationship between the infectivity and high transmissibility with low virulence has contributed to the ongoing global public health crisis, and the emergence of more transmissible and potentially more pathogenic variants of concern (VOCs). Longitudinal cross-sectional studies have indicated only a fraction of asymptomatic to mild and moderate SARS-CoV-2 natural infections of humans results in detectable neutralizing antibodies months after recovery (*13-15*) and current evidence suggests that convalescent immunity may only provide transient protection (*16-18*). Furthermore, the degree of immune evasion witnessed *in vitro* by spike (S) protein variants and the increasing occurrence of reinfection (*19-23*) has raised the question whether previous infection or vaccine-derived immunity can provide heterologous protection against further transmission and limit additional mutation of circulating VOC, even while protection against severe disease and mortality has been sustained to date.

Some VOCs show increased transmissibility, cause more severe disease than original strains and demonstrate greater immune evasion to neutralizing antibodies as they harbor mutations at sites facilitating viral fusion (*24, 25*) and at critical epitopes in the N-terminal (NTD) and receptor binding domains (RBD). The B.1.1.7 (Alpha), B.1.351 (Beta) and P.1 (Gamma) variants contain the same set of mutations, K417T, E484K, and N501Y, while the B.1.617 (Delta) carries L452R and T478K mutations, all of which are associated with enhanced infectivity and reduced serum neutralization (*26-28*). These antigenic changes have abolished potent neutralizing epitopes targeted by monoclonal antibody therapeutics (*29*) and antibodies elicited by mRNA vaccination (*30*). Natural reinfection with the VOC strains after original strain infection, and *vice versa*, has been documented (*20, 31-34*). Despite this, cross-neutralization of convalescent sera from patients infected with non-variants and mRNA immunized individuals suggests that the parental spike (S) protein in current vaccines affords some degree of protection (*35-37*).

Phase 1-3 clinical trials of novel mRNA (Moderna mRNA-1273 and Pfizer-BioNTech BNT162b2) and viral-vector (Johnson & Johnson-Janssen Ad26.COV2.S) platform, utilizing the original Wuhan-Hu-1 S protein (*38-40*), have shown these vaccines to be highly protective with 94-95% and 67% efficacy against COVID-19 disease, respectively (*41-43*), and they all elicit uniformly high protection against severe disease/death (i.e., >85%). Post-hoc sequencing during clinical trials of these and other vaccine platforms in variant-dominant regions have revealed a slightly lower efficacy against B.1.1.7 and B.1.351 variants, albeit usually remaining above the 50% protection threshold (*44, 45*). Attempts at predicting protection based on immune correlates, such as cross-neutralizing antibodies, have shown that vaccinated individuals had a 2 to 5-fold reduction in titers against current VOCs as IgG levels wane, which are nonetheless believed to confer a degree of resistance against infection (*46-49*). However, these samples were taken during the peak period of the humoral response and may not be accurate predictors of protection months onwards when the likelihood of infection is greater. Thus, understanding the decay in antibody responses months after the final boost is an essential endpoint in vaccine development.

Evaluating a delayed vaccine response in non-human primate (NHP) models can provide valuable foresight into the relationship between waning immunity and protection, especially against VOCs. In this study, we describe the immunogenicity and protective efficacy of a recombinant pre-fusion spike subunit (based on the reference SARS-CoV-2 strain, Wuhan-Hu- 1) adjuvanted with liquid or lyophilized (dry) CoVaccine HT™, a squalane-in-water nanoemulsion adjuvant containing immunostimulatory sucrose fatty acid sulfate esters (*50*), in cynomolgus macaques. Other investigators have shown that infection with SARS-CoV-2 (prototype strain) produced COVID-19-like disease in this species and reflects similar viral shedding kinetics and lung pathology as human infections (*51, 52*). We have previously demonstrated that our vaccine candidate elicits a broad-spectrum IgG response including high neutralizing antibody (NtAb) titers against the prototypic SARS-CoV-2 and VOCs, specifically B.1.351 and P.1, and an *in vitro* antigen-specific IFN-γ secreting response from immune splenocytes taken from Swiss Webster mice (*53*). To elaborate on our understanding of the elicited immunological response, we assessed the vaccine efficacy using a delayed challenge scheme with the P.1 VOC to delineate how the maturation and/or decay of the humoral response affects protection from a heterologous SARS-CoV-2 strain. Neutralizing antibody responses to the vaccine-matched WA1/2020 strain, and VOCs, B.1.351, P.1 and B.1.617 were also determined just prior to challenge and 14 days post-challenge (to assess anamnestic responses). Furthermore, we characterize the viral kinetics and histopathological changes post-challenge in the lower and upper respiratory tracts of control (non-vaccinated, challenged), and protected NHPs. To our knowledge, this is the first published account of P.1 VOC challenge in NHPs.

## RESULTS

### Spike antigen with either liquid or lyophilized CoVaccine HT™ elicits a durable humoral response

Twelve Cynomolgus macaques were assigned into four groups and received two immunizations with 5 (Group A) or 25 (Group B and C) μg liquid prefusion spike trimer (S) antigen formulated with either lyophilized (Group A and C) or liquid (Group B) CoVaccine HT™ adjuvant, or one dose of a co-lyophilized CoVaccine HT™-adjuvanted control containing an unrelated viral glycoprotein antigen (Group D) (Fig. 1A). The two doses were administered within a three-week interval and all NHPs were challenged with the P.1 isolate 12 weeks post-boost (∼ 3 months, study week 15). Wuhan-Hu-1 S- and RBD-specific IgG titers were measured by a multiplexed microsphere immunoassay (MIA) using insect cell expressed antigens coupled onto spectrally distinct, magnetic beads as described previously (*53*). Serum S-specific IgG concentrations were interpolated using a standard curve generated from S-specific human IgG purified from vaccinated individuals (Fig. 1B). RBD-specific IgG titers were read out as median fluorescence intensity (MFI) (Fig. S1). All NHP immunized with the adjuvanted S at both antigen doses seroconverted after the prime (week 3) with S-specific antibodies in the range of 20 to 70 μg/mL, and peak serum IgG concentrations detected two weeks after the boost (week 5) in the range of 70 to 753 μg/mL. Macaques given a 25 μg dose of S demonstrated a greater IgG response to the antigen compared to those receiving 5 μg. RBD-specific IgG titers followed a similar trend. As expected, animals in group D receiving an unrelated antigen did not develop any detectable S-specific IgG during this phase of the study. S-specific IgG remained detectable 12 weeks after the boost (week 15) although IgG concentrations dropped 3.0 to 9.9-fold relative to the prior peak titer.

**Figure 1.**
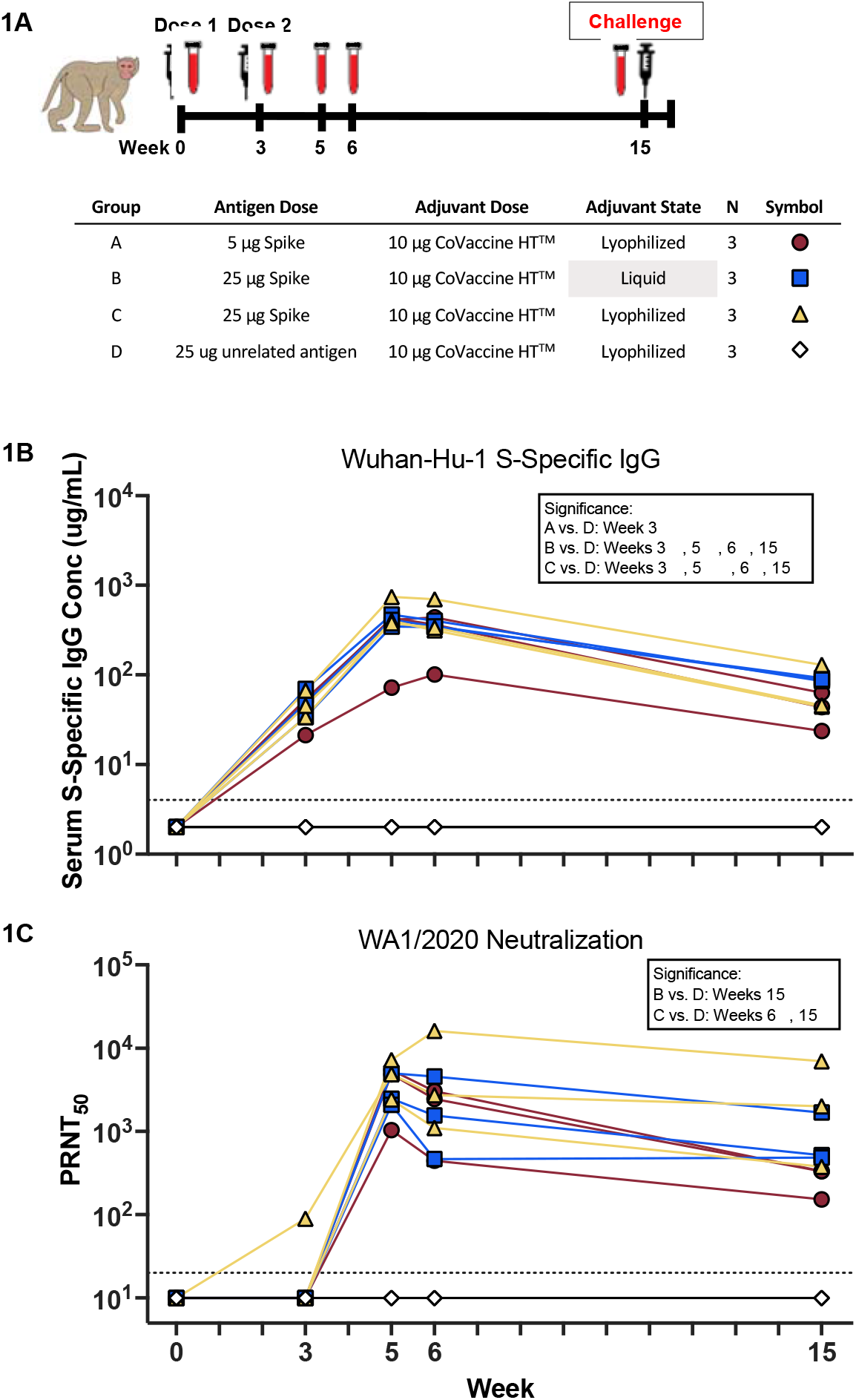
Vaccine Scheme, IgG and Neutralizing Antibody Kinetics. (A) 12 cynomolgus macaques were separated into 4 groups and given either 5 or 25 μg S protein formulated with either liquid or reconstituted, lyophilized CoVaccine HT™ adjuvant. Two doses were administered IM 3 weeks apart and sera was collected at indicated time points. Macaques were challenged IN and IT with a total of 1×10^6^ TCID_50_ of the SARS-CoV-2 P.1 strain. (B) Serum Wuhan-Hu-1 S-specific IgG kinetics measured using a MIA with purified, human S-specific IgG standards to estimate serum concentration. Dashed line indicates the limit of quantification (LOQ, ≤4 μg/mL). Individual values falling below the LOQ were set to ½ LOQ. Significance was calculated using a one-way ANOVA followed by a Dunnett’s Multiple Comparison to Group D at each time point. (C) WA1/2020 neutralizing antibody kinetics were measured using a WT SARS-CoV-2 PRNT assay. Curve-fitted PRNT_50_ titers were calculated using a sigmoidal dose response curve. Dashed line indicates the limit of detection (LOD, ≤1:20). Individual values falling below the LOD were set to ½ LOD. Significance was calculated using a Kruskal-Wallis Test followed by a Dunn’s Multiple Comparison. (*p≤0.05, **p≤0.01).

The titer of NtAb against the prototype WA1/2020 strain was determined using a standard PRNT with wild-type virus (Fig. 1C). NtAb from one animal in Group C was detectable after the prime (week 3). All S-immunized NHPs developed a potent neutralizing response, peaking two weeks after the boost (week 5) and generally remaining stable one week later (week 6). The group receiving 5 μg of the antigen showed the greatest variability. Nearly all immunized NHPs maintained neutralization capacity, with PRNT_50_ greater than 1:150 dilution, 12 weeks after the final immunization (week 15). NtAbs from groups receiving 25 μg S either remained stable or decreased up to 3.0-fold, while the group receiving 5 μg S declined ∼2.9-9.2-fold. A similar reduction in serum neutralization was also verified using a surrogate rVSV-SARS-CoV-2 S PRNT assay (Fig. S2).

### A recombinant subunit vaccine induces stable neutralizing antibodies against VOCs

Circulating SARS-CoV-2 VOCs can evade vaccine-induced antibody responses and are associated with breakthrough infections in those fully immunized, especially as antibody titers wane (*54-60*). To determine whether an adjuvanted, prototypic Wuhan-Hu-1 S subunit can generate durable cross-variant neutralizing antibodies, PRNTs using WT B.1.351, P.1, and B.1.617 isolates were determined with sera collected at the time of peak neutralization and 12 months after the final immunization, a timepoint at which antibody titers are expected to have waned. At week 6, potent neutralization of the B.1.351, P.1 and B.1.617 VOC was detected, although 10.7-, 10.7- and 5.7-fold lower, respectively, compared to WA1/2020 in all vaccine groups (Fig. 2A). As antibody titers waned 10 weeks later (study week 15) the gap between neutralization titers against the WA1/2020 strain to the VOC isolates narrowed to a 4.5-, 6.3- and 5.1-fold difference (Fig. 2B) driven by a proportionally larger decline in WA1/2020-specific neutralizing activity. At this later time point, all S-immunized macaques, except for a single macaque in Group A, maintained a detectable PRNT_50_ titer greater than 1:40 dilution. A few macaques in Groups B and C demonstrated greater VOC neutralization at week 15 compared to week 6, suggesting a refinement or “maturation” of the humoral response towards neutralizing epitopes during this interval.

**Figure 2.**
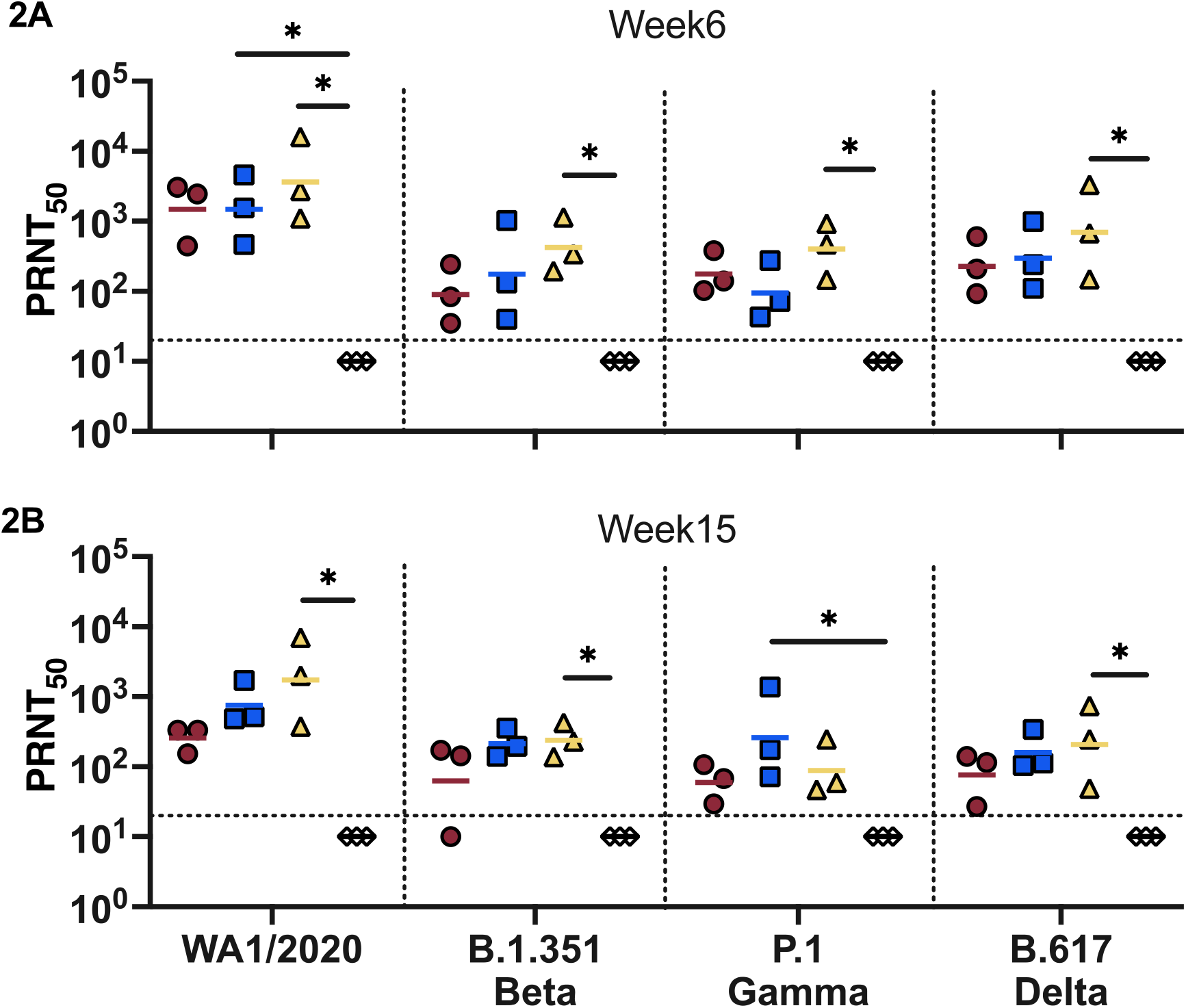
Cross-Neutralizing Antibody Titers. Neutralizing antibody titers at (A) week 6 and (B) week 15 were measured using a WT SARS-CoV-2 PRNT assay with the WA1/2020, B.1.351 (Beta), P.1 (Gamma), B.1.617 (Delta) VOC. Curve-fitted PRNT_50_ titers were calculated using a sigmoidal dose response curve. Dashed line indicates the LOD (≤1:20). Individual values falling below the LOD were set to ½ LOD. Significance was calculated using a Kruskal-Wallis Test followed by a Dunn’s Multiple Comparison to Group D at each time point. Horizontal bars mark the GMT of each group (*p≤0.05).

### Immunization with recombinant subunits reduces viral burden from delayed P.1 (Gamma) challenge

The characterization of the prototype strain and B.1.351 variant infections in NHP models has been described previously (*51, 52, 61-63*). Here we characterize the clinical signs and histological events of a P.1 VOC infection in cynomolgus macaques and determine whether a prefusion spike subunit formulated with CoVaccine HT™ is effective at reducing viral load against a circulating VOC after a 12-week interval following the final immunization. All NHPs were challenged with a total of 1×10^6^ TCID_50_ of the P.1 isolate using simultaneous intranasal and intratracheal inoculation routes. None of the NHPs challenged developed visible clinical signs of respiratory disease throughout the study, consistent with WA1/2020 strain infected cynomolgus macaques (*51*). Bronchioalveolar lavages (BAL) from the lower respiratory tract, and nasal and oral swabs (NS and OS respectively) from the upper respiratory tract were collected at days 2, 4, 7, 10 and 14 after challenge to detect infectious virus, genomic RNA or signs of viral replication. High levels of infectious virus, GMT Log_10_ TCID_50_ titer of 5.8 and 4.8, were recovered two days post-challenge from the NS and BAL, respectively, from control macaques (Group D, Fig. 3A & 3D and S3A-B). Infectious virus continued to be detectable in both anatomic sites until at least day 7 post-challenge before becoming undetectable on day 10. All S-immunized NHPs presented TCID_50_ titers at least 1-2 Log_10_ lower than control macaques in both locations at all timepoints throughout the study, indicating protection conferred by both 5 and 25 μg S formulations of the subunit vaccine. No infectious virus could be cultured from the BAL in S-immunized macaques by day 7. Infectious virus from the NS was recoverable from immunized macaques throughout the study period and was variable between groups, however these macaques on average had TCID_50_ at least 1-2 Log_10_ below those of control animals.

**Figure 3.**
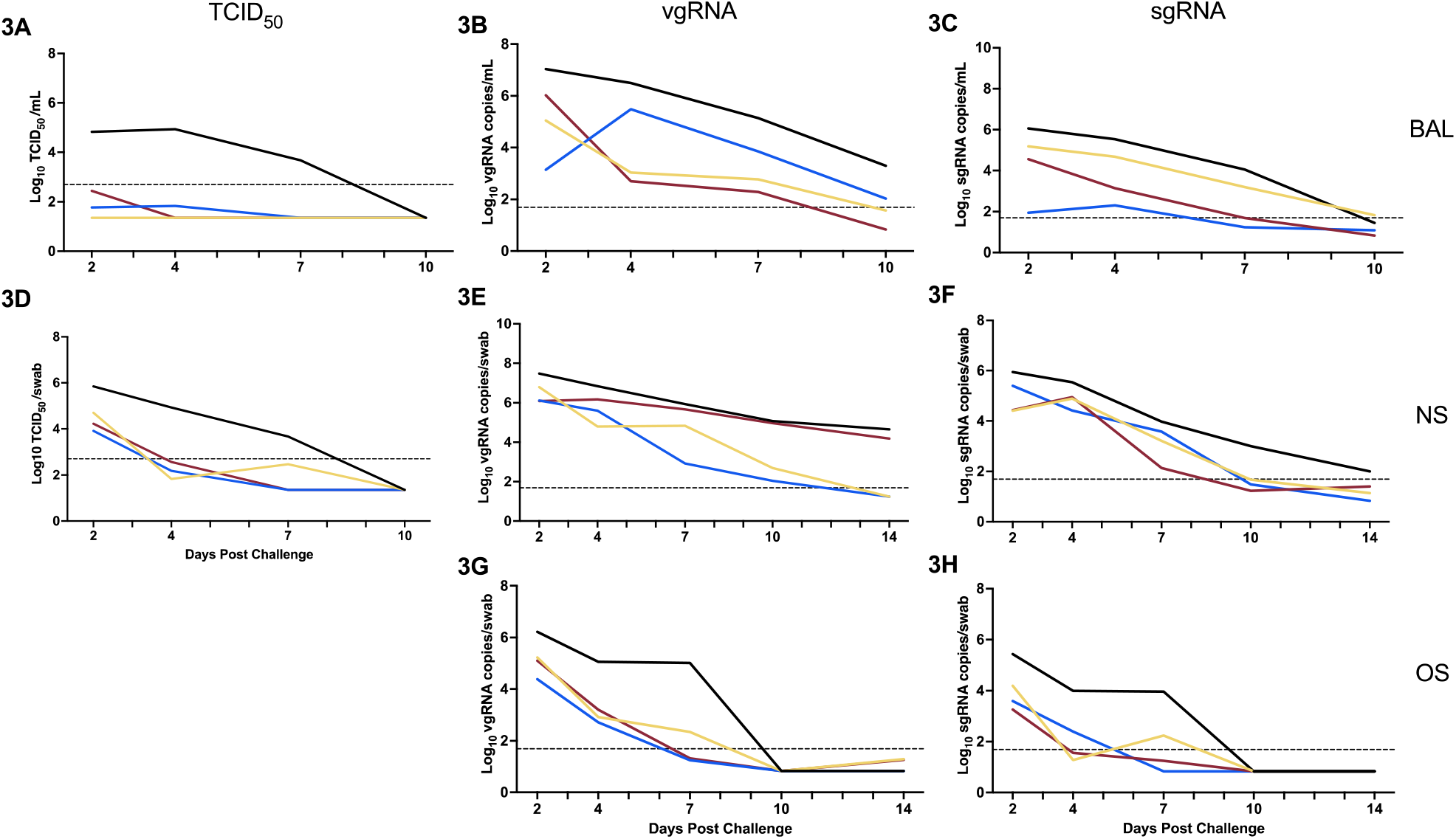
Viral Load Kinetics after P.1 Challenge. GMT of the (A & D) TCID_50_ titers; (B, E, G) viral genomic; and (C, F, H) subgenomic N RNA copies from the (A, B, C) bronchoalveolar lavage (BAL); (D, E, F) nasal swab (NS); and (G, H) oral swab (OS) collected at the time points indicated. Dashed line indicates the assay-specific LOD. Individual values falling below the LOD were set to ½ LOD.

Quantitative RT-PCR was used as an additional measurement for viral load. We detected the presence of viral RNA and viral replication throughout the 14-day study period in both the lower (Fig 3B-C, S4A-C and S5A-C) and upper (Fig 3E-F, S4D-F and S5D-F) respiratory airways and from the oral cavity (Fig 3G-H, S4G-I and S5G-I). Quantification of viral RNA showed that both 5 and 25 μg immunization had reduced the viral load by approximately 1-2 Log_10_ RNA copies/mL (or swab) over Group D macaques, corroborating the TCID_50_ results. Virus replication in the oral cavity was undetectable at least 3 days earlier in most S-immunized macaques than in the control group, suggesting a reduced potential for transmission. High levels of viral RNA and replication, between 5-6 Log_10_ vgRNA and sgRNA copies/mL, respectively, were observed in these control macaques before gradually resolving at day 10. Altogether, reduced viral loads at all tested anatomic locations in Groups A, B and C macaques indicate a degree of durable protection from P.1 VOC infection provided by an adjuvanted protein subunit vaccine.

### High dose spike antigen reduces lung histopathology caused by P.1 (Gamma) challenge

Also consistent with previous prototype virus infection studies in cynomolgus macaques (*51, 52*), P.1 challenge did not cause changes in weight, temperature nor demonstrate any observable signs of respiratory disease. To evaluate vaccine efficacy against COVID-19-like pathology, a section slide from each lung lobe and the bronchi was prepared from each macaque and stained with hematoxylin and eosin to survey for histopathological changes indicative of acute lung injury. The histopathological scoring system based on the presence/severity of edema, intraalveolar and interstitial inflammation, perivascular lymphocytic cuffing, and increased bronchiolar-associated lymphoid tissue (BALT) is outlined in Table S1. A cumulative average score was determined for each macaque based on the evaluation of these five characteristics per section for a total of 30 scores (Fig 4A). Significant differences in the cumulative average scores for each vaccine formulation vs. controls were determined using a one-way ANOVA followed by a Dunnett’s multiple comparison test. Macaques in the control Group D appeared to have developed mild to moderate respiratory disease. The lower cumulative GMT scores of Groups B and C, immunized with 25 μg S, suggests that these animals, except for a single macaque, were completely protected from lung pathology while all Group A (immunized with 5 μg S) macaques appeared to have developed mild disease despite exhibiting lower viral loads. Week 15 WA1/2020 NtAb levels were determined to be inversely correlated, although weakly, with histopathology scores using a Spearman’s correlation test (r=- 0.682, p=0.0178) (Fig. 4B). Week 15 pre-challenge neutralizing antibody levels were lower in animals with higher histopathology scores, a trend seen with breakthrough infections in fully immunized individuals (*64*). The moderate disease seen in the control animals is demonstrated in Fig. 5A, where a representative, low-power magnification shows a distribution of lymphocytic perivascular cuffing and clusters of intraalveolar macrophages. In contrast, immunized animals (especially those receiving the higher antigen dose), appeared to have milder disease without these pathological changes (Fig. 5B). Macaques developing mild to moderate disease exhibited absent to moderate, focal and multifocal edema and perivascular cuffing with marked lymphocytic infiltration (Figs. 6A-D). Syncytia of intraalveolar multinucleated giant cells surrounded by acute and eosinophilic inflammatory infiltrate (Fig. 6A.) or increased intraalveolar macrophages with interstitial lymphocytic inflammation, in addition to alveolar septal thickening and complete perivascular cuffing can be seen in a moderately diseased, control macaque (Fig. 6B-C). Proteinaceous edema fluid filling alveolar spaces was also observed in unprotected macaques (Fig. 6D). While these histopathological changes were also noticed in protected macaques, the findings appeared less severe (Fig. 6E-F). A section from a protected macaque revealed a few intraalveolar multinucleated giant cells without an increase in intraalveolar macrophages, lymphocytic vascular cuffing, or interstitial inflammation (Fig. 6E). Similarly, only partial lymphocytic perivascular cuffing with adjacent alveolar epithelial hyperplasia was noticed without an increased intraepithelial macrophage infiltrate and normal alveolar septal thickness in immunized macaques receiving 25 μg of S (Fig. 6F).

**Figure 4.**
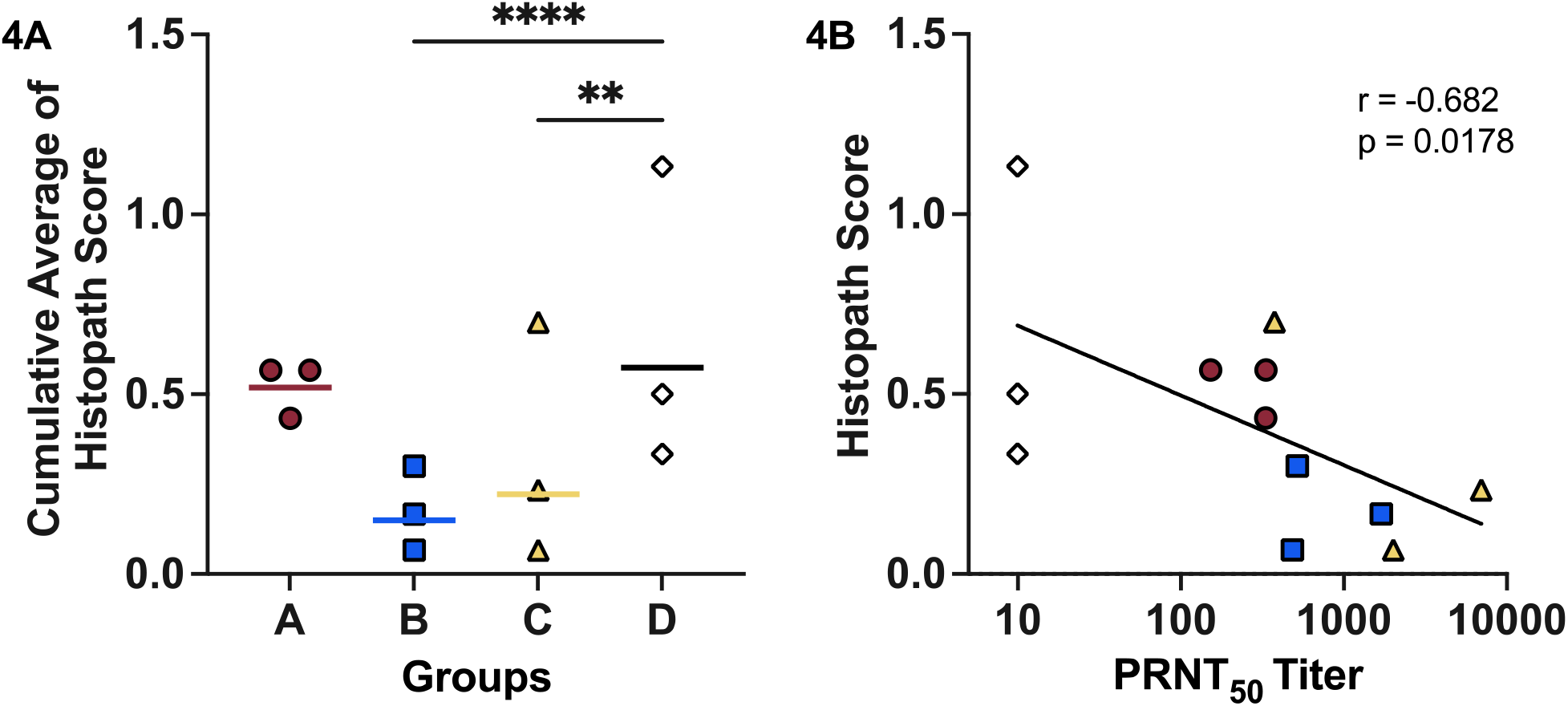
Histopathology Score and Correlation with Neutralizing Antibodies. (A) Cumulative average histopathology score was determined by averaging the scores of six sections cut from each lobe and the bronchi. The presence and severity of edema, intra-alveolar inflammation, perivascular cuffing, increased BALT and interstitial inflammation was determined and assigned scores ranging from 0-3. Significant differences in lung histopathology were calculated using a one-way ANOVA followed by a Dunnett’s multiple comparison to Group D with the scores from each section (30 scores per macaque) as replicates. Horizontal bars mark the GMT of each group. (**p≤0.01, ****p≤0.0001) (B) Correlation between histopathology scores and WA1/2020-specific PRNT_50_ titers were determine using a Spearman’s correlation test.

**Figure 5.**
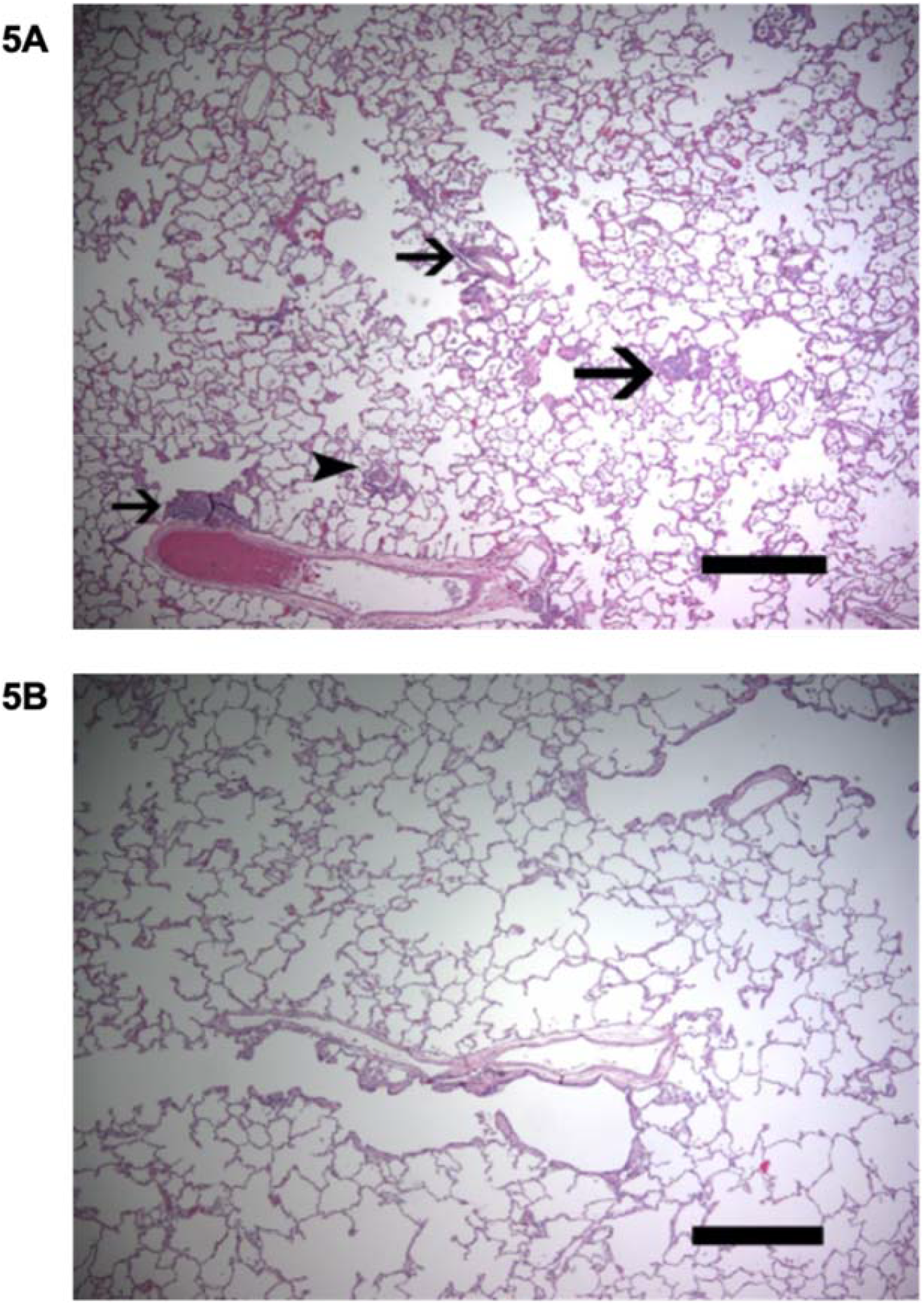
Histopathology observed in Hematoxylin and Eosin-Stained Lung Sections of Control Cynomolgus Macaques Compared to Immunized Macaques after P.1 challenge. (A) Representative low power view of a lung section showing distribution of lymphocytic perivascular cuffing (arrows) and clusters of intraalveolar macrophages (arrowhead) in an unprotected macaque. (B) Representative low power view of a lung section from an immunized macaque which did not exhibit pathologic changes. Scale Bar = 500 microns.

**Figure 6.**
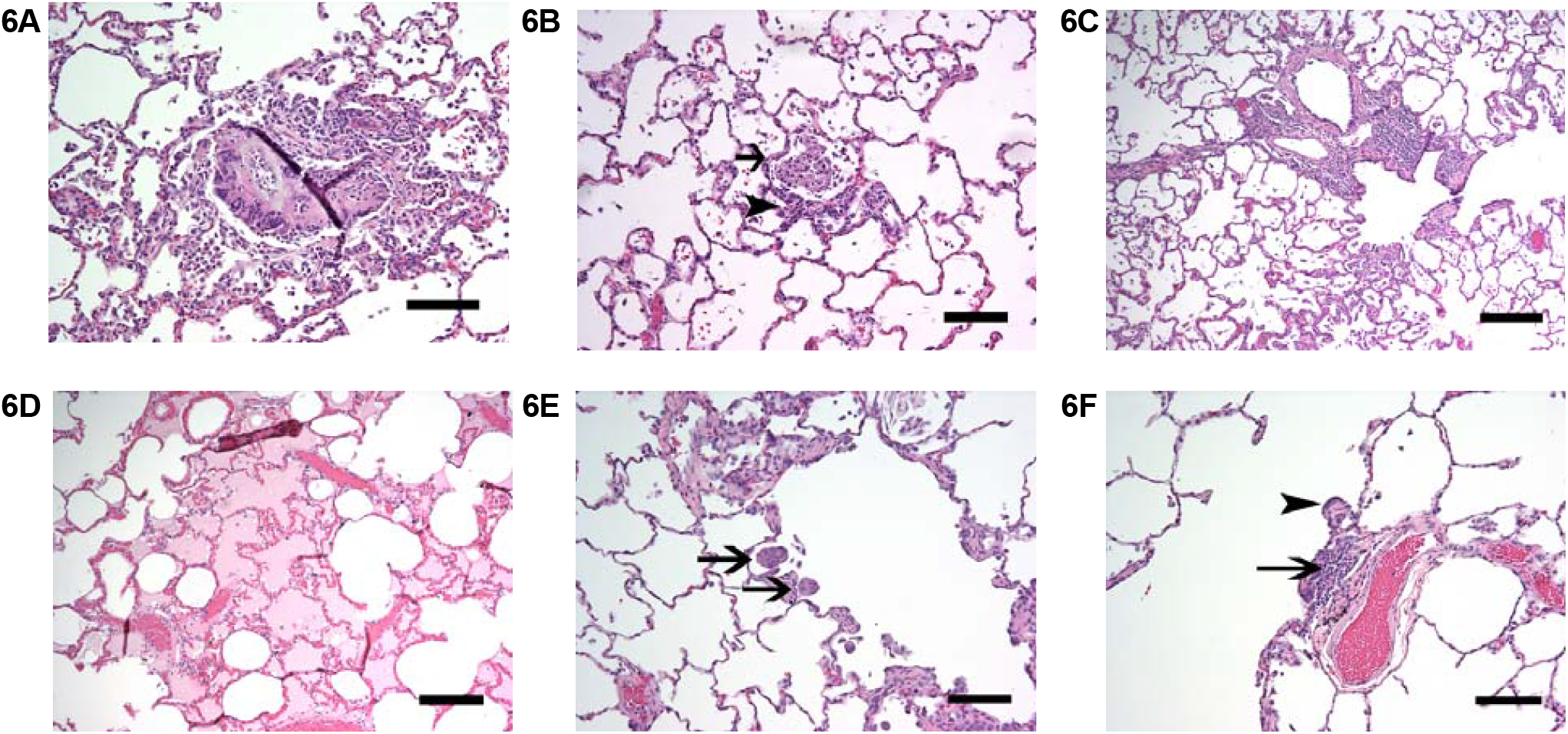
Histopathology Observed in Control Cynomolgus Macaques Compared to that Observed in Immunized Macaques after P.1 Challenge. (A-D) unprotected macaques. (A) A syncytium of intraalveolar multinucleated giant cells is seen surrounded by an acute and eosinophilic inflammatory infiltrate. (B) Increased intraalveolar macrophages and an intraalveolar multinucleated giant cell accompanied by interstitial lymphocytic inflammation and thickening. (C) Complete lymphocytic perivascular cuffing with increased intra-alveolar macrophages. (D) Proteinaceous edema fluid in alveolar spaces. (E-F) Protected macaques. (E) Intraalveolar multinucleated giant cells (arrows) without increased intraalveolar macrophages, lymphocytic vascular cuffing or interstitial inflammation. (F) Partial lymphocytic perivascular cuff (arrow) with adjacent alveolar epithelial hyperplasia (arrowhead) without alveolar septal thickening. Scale bar = 10 microns.

### P.1 challenge recalls a broadly neutralizing anamnestic response and a variant-specific primary neutralizing response in naïve macaques

To better understand the immunological basis of events happening after late challenge, and to determine if P.1 challenge triggered an anamnestic response in immunized NHPs, post-challenge S-specific IgG concentrations and PRNT_50_ titers were measured from sera collected upon necropsy at day 14. Late P.1 challenge significantly boosted serum IgG concentrations to levels attained at the post-immunization peak at Week 6, with less variance between animals, suggesting that a recall response was triggered by the heterologous infection (Fig. 7A). Vaccine-matched (WA1/2020) and VOC neutralizing (i.e., B.1.351, P.1 and B.1.617) antibody levels were also all significantly boosted to levels attaining or surpassing the peak levels generated through initial immunization by 5-fold and upwards of 20-fold respectively (Fig. 7B and S2). Unsurprisingly, the greatest increase from pre-challenge neutralization was against the P.1 challenge isolate. Group A showed slightly higher P.1 NtAbs compared to Group B and C macaques and was the only group to have significantly higher titers across different variants. The lower degree of histopathology seen in Groups B and C may indicate that 25 μg S with either form of CoVaccine HT™ provides greater long-term protection. In Group D control macaques, P.1 challenge generated low levels of NtAbs compared to immunization alone, and it was only cross-reactive with the B.1.351 isolate and not to WA1/2020 nor B.1.617 isolates. This suggests that primary infection with one isolate of SARS-CoV-2 does not generate broad immunity against other variants in contrast to vaccine derived immunity, which is more broadly cross-reactive with VOCs, even months after peak immunity is observed.

**Figure 7.**
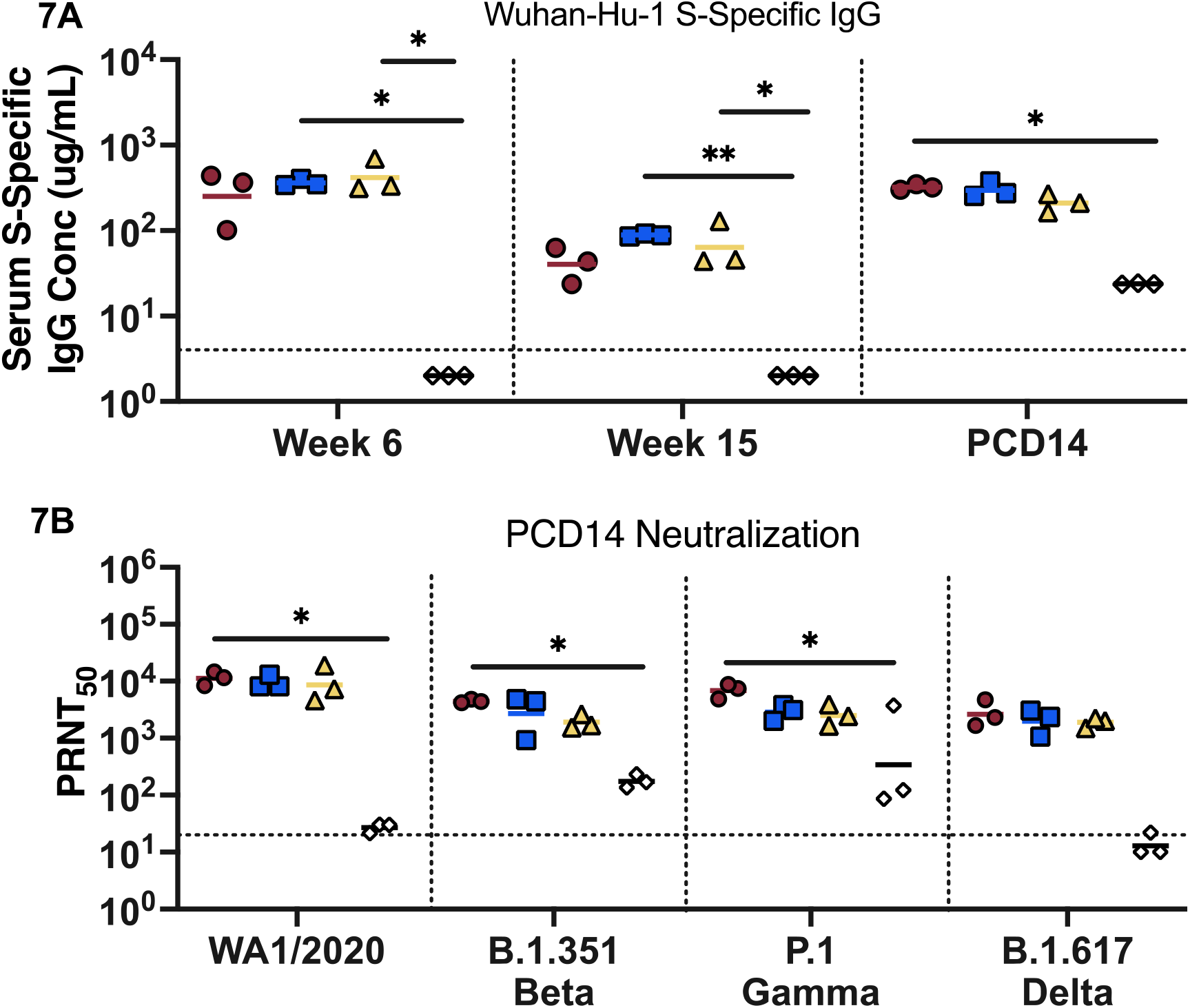
Post-Challenge Anamnestic Response. (A) Vaccine-matched S-specific IgG conc. from sera collected at week 6, week 15 and at 14-days post-challenge, measured by MIA. IgG concentrations were estimated using a human S-specific IgG standard as described previously. Horizonal bars mark the GMT for each group. Dashed line indicates the LOQ (≤4 μg/mL) Individual values falling below the LOQ were set to ½ LOQ (B) Neutralizing antibodies 14 days after P.1 challenge were measured using a WT SARS-CoV-2 PRNT assay with the WA1/2020, B.1.351 (Beta), P.1 (Gamma), B.1.617 (Delta) VOC. Curve-fitted PRNT_50_ titers were calculated using a sigmoidal dose response curve. Dashed line indicates the LOD (≤1:20). Individual values falling below the LOD were set to ½ LOD. Significance was calculated using a Kruskal-Wallis Test followed by a Dunn’s Multiple Comparison to Group D at each time point. (*p≤0.05, **p≤0.01).

## DISCUSSION

Understanding the durability and breadth of vaccine-generated immunity is critical for predicting long-term protection during the COVID-19 pandemic and for informing policy on strategic resource deployment to facilitate equitable vaccine access. Diversifying the type of vaccines currently used beyond mRNA and viral vectors to include other vaccine platforms, such as protein subunit vaccines, can bolster global availability by mobilizing more thermostable vaccines to resource-poor areas, or overcoming anti-vector immunity and adverse effects, by using this platform as a booster to restore pre-existing natural or vaccine-induced immunity (*65-67*).

We have demonstrated that two doses of a subunit vaccine consisting of a prefusion S trimer (Wuhan-Hu-1) formulated with liquid or lyophilized CoVaccine HT™ reduces viral load and provides sufficient, durable, cross-variant protection from mild to moderate disease lasting at least 3 months after the final boost. This supports previous findings in rhesus macaques that S trimer-based subunit protein vaccines are highly immunogenic and generate long-lasting, robust antibody responses (*68*). Furthermore, using a two-step purification method of immunoaffinity chromatography followed by a size-exclusion polishing step results in a homogenous antigen composition and reduces host-cell carryover plaguing conventional purification strategies for protein subunits (*69*). Also promising is the prospect that the adjuvant, CoVaccine HT™, can be lyophilized, and reconstituted simply with water for injection and still retain functionality. This technology has been previously used to develop mono- and multivalent filovirus vaccine formulations. Those studies demonstrated that single-vial, lyophilized formulations preserved the higher order antigen structure and the biophysical properties of the adjuvant (*70, 71*). Robust levels of S-specific IgG and homologous, as well as B.1.351, P.1 and B.1.617 cross-neutralizing antibodies, were detected throughout the extended study period and inversely correlate with viral load and lung damage. This report supports growing evidence that the Wuhan-Hu-1 S peptide sequence encoded by highly effective mRNA, viral vectored, and subunit protein vaccines, generates immunity that affords protection against circulating VOCs (*45, 46, 70-73*), even months after the final vaccine dose when antibody titers are waning. Furthermore, we noticed a few macaques receiving the higher antigen dose had developed increasingly potent cross-variant neutralizing titers immediately prior to viral challenge which were greater than titers observed during the peak humoral response. This suggests that a potentially remaining antigen depot or persistence of APC’s may foster continued accumulation of somatic mutations and affinity maturation in memory B cells (*74, 75*) beyond the initial vaccination phase.

This study is also the first to describe the course of infection and histopathology of a SARS-CoV-2 P.1 infection in a NHP model. Consistent with what was described in WA1/2020 infected cynomolgus macaques (*51, 52*), macaques challenged with the P.1 isolate did not develop elevated temperature or decreased weight, nor did they show observable signs of respiratory disease. Viral load in the upper and lower respiratory tract peaked early during infection two days after challenge and gradually decreased to undetectable sgRNA levels by day 14 in unprotected animals, similar to viral kinetics described in another study using the same inoculating dose (1×10^6^ TCID_50_), but with an early pandemic isolate (*52*). Although P.1 is estimated to be upwards of 2.4-fold more transmissible (*76*), it is unclear whether the higher viral load observed during peak P.1 replication in this study compared to time-matched reports for a non-variant isolate is a characteristic of P.1 replication or a discrepancy between readouts from different RT-PCR procedures. However, we show that immunization with both 5 and 25 μg doses of S effectively reduces viral load in both upper and lower respiratory tracts implying a lower likelihood of viral transmission even in mildly symptomatic macaques. Histopathological examination of lung and bronchial sections in unprotected macaques confirmed mild to moderate disease, which was abrogated in immunized individuals. COVID-19 like disease observed in this model consisted of increased intraalveolar and interstitial infiltration, as well as giant syncytial cells, similar to previous observations (*51, 77, 78*). While both high and low doses of S antigen generate statistically negligible differences in humoral responses during peak immunity, it is clear from our findings that higher antigen doses, at least with CoVaccine HT™ produces a more durable and protective response at this later timepoint.

Late P.1 challenge induced an anamnestic response in all immunized macaques that restored S-specific antibody titers to peak serum concentrations seen shortly after the final boost, and significantly enhanced vaccine-matched and VOC-neutralizing antibody titers. In most macaques receiving 25 μg S, the immune recall occurred with little to no disease. The durability and cross-variant neutralizing nature of immune responses generated by mRNA and viral vectors has been documented up to six months or later (*79-82*). However, breakthrough infections, particularly with variant strains, have been reported even in the presence of detectable and high-level NtAbs (*54, 56, 59, 83*). Anamnestic responses in fully immunized NHPs have been characterized previously with homologous and B.1.351 variant challenge and agree with our observations that viral challenge boosts functional antibody levels. Our analysis shows that viral challenge with a heterologous strain not only rapidly boosts homologous and challenge strain specific NtAbs, but also NtAbs against the unencountered variants, B.1.351 and B.1.617, underpinning the broad-spectrum potential of subunit protein vaccines. In contrast, control macaques, receiving no S antigen, developed moderate levels of P.1 NtAbs that are only cross-neutralizing with B.1.351, and barely neutralize WA1/2020 and B.1.617, suggesting that natural infection with one strain of SARS-CoV-2 may confer only limited protection against other VOCs. Of course, further affinity maturation could not be observed here as the study ended 14 days after challenge.

The small size of our treatment groups (n=3) is a limitation of our study and may therefore not provide enough statistical power to strongly correlate antibody concentrations to viral load or histopathology score. Furthermore, we cannot directly compare the outcome of our study to other studies that evaluate vaccine efficacy soon after the final booster when immunity is greatest, as vaccine efficacy is known to decline over time. Likewise, we could not benchmark the efficacy of our vaccine formulation when the immunity is greatest at week 5, however, our late challenge scheme accurately reflects the current urgent need for decisions about timing and candidates for possible booster vaccinations during the pandemic situation as it provides useful information regarding the real-life durability and breadth of vaccine protection. It furthermore may shed light into the utility of protein vaccines to serve as prime or boost in combination with other vaccines.

In conclusion, we show that a two-dose regimen of a prefusion-stabilized trimeric S subunit protein vaccine formulated with lyophilizable CoVaccine HT™ adjuvant reduces viral burden and high antigen doses can confer durable cross-variant immunity. Future efforts will therefore focus on developing a thermostabilized vaccine formulation in a single-vial presentation, potentially enabling facile worldwide distribution.

## MATERIALS AND METHODS

### Ethical Statement

The investigators adhered fully to the “Guide for the Care and Use of Laboratory Animals” by the Committee on Care of Laboratory Animal Resources Commission on Life Sciences, National Research Council. Cynomolgus macaques (*Macaca fascicularis*) were housed at BIOQUAL Inc. (Rockville, MD). All macaque experiments were reviewed and approved by BIOQUAL’s Animal Care and Use Committee. BIOQUAL Inc. is accredited by the American Association for Accreditation of Laboratory Animal Care (AAALAC).

### Study Design

Cynomolgus macaque studies were performed using three adjuvanted vaccine formulations and an adjuvanted control using unrelated antigens: A) 5 μg of SARS-CoV-2 S protein with 10 mg lyophilized CoVaccine HT™, B) 25 μg S protein with 10 mg liquid CoVaccine HT™, C) 25 μg S protein with 10 mg lyophilized CoVaccine HT™, and D) 25 μg of Ebola Virus (EBOV) glycoprotein with 10 mg CoVaccine HT™, co-lyophilized in one vial. CoVaccine HT™ (BTG International Ltd, London, United Kingdom) was lyophilized as previously described (*71*) and reconstituted with sterile PBS before mixing with the antigen. Each group consisted of both male and female cynomolgus macaques (*n* = 3 for each formulation) weighing between 2.8 and 8.2 kg. Cynomolgus macaques of groups A-C were immunized intramuscularly (IM) in both deltoids (split dose) at weeks 0 and 3, control animals were immunized only once at week 0. Pre-challenge sera were collected at weeks 0, 3, 5, 6, 15. Challenge at week 15 with SARS-CoV-2 P.1 variant, hCoV-19/Japan/TY7-501/2021, TY7-501 (BIOQUAL-generated stock [lot no. 031921-1215] in Calu-3 cells from seed stock no.TY7-501 was performed with an inoculum dose of 5×10^5^ TCID_50_/mL administered to each animal in volumes of 1 mL by intratracheal and intranasal injection at each site. Nasal and oral swabs (NS and OS respectively) were collected on days 2, 4, 7, 10 and 14 after challenge. Bronchoalveolar lavage (BAL) samples were collected on days 2, 4, 7 and 10 after challenge. Necropsies and lung tissue collection were performed at the endpoint of the study at day 14 post-challenge. Samples were immediately processed and subsequently stored at −80°C prior to analysis. S-specific serum IgG concentrations were determined using a microsphere immunoassay (MIA; see below). NtAbs were measured using wild-type and rVSV-SARS-CoV-2 S PRNTs (see below). Viral load in the NS, OS and BAL was measured using plaque assays and quantification of genomic and subgenomic N transcript RNA (see below). Lung and bronchi tissues were processed, and H&E stained to delineate histopathological signs of disease in each animal.

### Recombinant protein expression and purification

Plasmids were generated to express the pre-fusion, protease-resistant, trimeric transmembrane (TM)-deleted spike (S) glycoprotein from SARS-CoV-2 as described previously (*53*). Modifications to the gene include the removal of the furin and S2’ cleavage site, the addition of 2 prolines between the heptad repeat 1 and central helix region, and a foldon trimerization domain. A stably transformed cell line was created by hygromycin B selection at 300 μg/mL. The cell line was scaled up to 1L using a WAVE bioreactor (Cytiva, Marlborough, MA) and induced with 200 μM CuSO_4_.

Recombinant S protein was purified from clarified cell culture supernatants by immunoaffinity chromatography (IAC) using the SARS-CoV-2 cross-reactive mAb CR3022 (provided by Mapp Biopharmaceutical) coupled to NHS-activated Sepharose at a concentration of 10 mg/mL. The antigen was eluted with a glycine buffer (pH 2) in tandem into a HiPrep 26/10 desalting column (Cytiva, Marlborough, MA) equilibrated with PBS. The oligomeric content was evaluated by size-exclusion chromatography using a HiLoad 16/600 column (GE Healthcare, Piscataway, NJ) equilibrated with PBS. The S protein eluted as a single peak and the final product migrated as two bands, corresponding to the monomer and trimer on SDS-PAGE, under denaturing conditions, and was reactive to CR3022 mAb on a western blot. Antigens were sterile filtered with a 0.22 μm syringe filter (Cytiva, Marlborough, MA) and stored at -80°C until use.

### Analysis of antibodies by multiplex microsphere immunoassay (MIA)

The IgG antibody titers in sera were measured using a multiplex microsphere-based immunoassay as described previously (*53, 84-86*). Spectrally distinct, magnetic MagPlex® microspheres (Luminex Corporation, Austin, TX) were coupled to purified S, RBD or bovine serum albumin (BSA). A mixture of the antigen-coupled beads was incubated with sera diluted with PBS+ 1% BSA and 0.02% Tween 20 (PBS-BT) at 1:5,000 or 1:10,000 in black-sided 96-well plates for 3 hours at 37°C with agitation. Bound IgG was detected using 1μg/mL red phycoerythrin (R-PE)-conjugated goat anti-human IgG antibodies (Jackson ImmunoResearch, Inc., West Grove, PA) and resuspended in MAGPIX® drive fluid before being analyzed on a MAGPIX® Instrument (Luminex Corporation, Austin, TX).

To determine S-specific IgG concentrations in the sera, the median fluorescence intensity (MFI) readouts of each sample was interpolated against a standard curve generated using purified human IgG at concentrations in the range of 7.44 to 1000 ng/mL. To produce the antibody standard, IgG was purified from pooled sera of COVID-19 vaccinated human volunteers using protein A affinity chromatography, followed by immunoaffinity chromatography (IAC) using NHS-Sepharose (Cytiva, Marlborough, MA) coupled with recombinant S to select for S-specific IgG. Purity was assessed using SDS-PAGE and antibody concentration was quantified using UV_280_ absorbance. The resulting MFI values were plotted against the Log_10_-transformed concentrations and fitted using a sigmoidal dose-response, variable slope model (GraphPad Prism, San Diego, CA). The resulting curves yielded r^2^ values > 0.99 with a well-defined top and bottom and the linear range of the curve. The experimental S-specific IgG concentrations in experimental samples were determined by interpolation on the standard curves, multiplied by the dilution factors and plotted as antibody concentrations (ng/mL).

### Recombinant vesicular stomatitis virus (rVSV) neutralization assay

Replication-competent rVSV expressing SARS-CoV-2 S protein (Wuhan-Hu-1) was generated as described previously (*87*) and the virus stocks were amplified in Vero E6 cells. For the plaque reduction neutralization test (PRNT), individual NHP serum samples were heat-inactivated at 56°C for 30 minutes. Six 3-fold serial dilutions of serum samples, starting at 1:40 dilution, were prepared and incubated with 100 plaque-forming units (PFU) of rVSV-SARS-CoV-2-S at 37°C for 1 hour. Antibody-virus complexes were added to Vero cell monolayers in 6-well plates and incubated at 37°C for another hour followed by addition of overlay media mixed with 1% agarose. 72 hours later, cells were fixed and stained with a solution containing 1% formaldehyde, 1% methanol, and 0.05% crystal violet overnight for plaque enumeration. Neutralization titers (PRNT_50_) were generated using a variable slope, nonlinear regression, with upper and lower constraints (100% and 0% neutralization, respectively), using Prism 9 (GraphPad Software, San Diego, CA).

### TCID_50_ and Wild-type SARS-CoV-2 virus PRNT_50_ assay

TCID_50_ and PRNT_50_ assays were performed in a biosafety level 3 facility at BIOQUAL, Inc. (Rockville, MD). The TCID_50_ assay was conducted by addition of 10-fold graded dilutions of samples to Vero TMPRSS2 cell monolayers. Serial dilutions were performed in the cell culture wells in quadruplicates. Positive (virus stock of known infectious titer in the assay) and negative (medium only) control wells were included in each assay set-up. The plates were incubated at 37oC, 5.0% CO2 for 4 days. The cell monolayers were visually inspected for CPE, i.e. complete destruction of the monolayer with cellular agglutination. The TCID_50_ value was calculated using the Read-Muench formula (*88*). For samples which had less than 3 CPE positive wells, the TCID_50_ could not be calculated using the Reed-Muench formula, and these samples were assigned a titer of below the limit of detection (i.e., <2.7 log10 TCID_50_/mL). For optimal assay performance, the TCID50 value of the positive control should test within 2-fold of the expected value.

To measure neutralization, sera from each NHP were diluted to 1:10 followed by a 3 fold-serial dilution. Diluted samples were then incubated with 30 plaque-forming units of wild-type SARS-CoV-2 USA-WA1/2020 (BEI NR-52281), B.1.351 (BEI NR-55282), or P.1. (BEI NR-54982) variants, in an equal volume of culture medium for 1 hour at 37°C. The serum-virus mixtures were added to a monolayer of confluent Vero E6 cells and incubated for one hour at 37°C in 5% CO_2_. Each well was then overlaid with culture medium containing 0.5% methylcellulose and incubated for 3 days at 37°C in 5% CO_2_. The plates were then fixed with methanol at -20°C for 30 minutes and stained with 0.2% crystal violet for 30 min at room temperature. PRNT_50_ titers were calculated using a variable slope, nonlinear regression, with upper and lower constraints (100% and 0% neutralization, respectively), on Prism 9 (Graphpad Software, San Diego, CA).

### Histopathology

NHP Lung tissue specimens from each lung lobe and bronchi were harvested at time of necropsy and preserved in 10% formalin before processing and parafilm embedded, fixed and stained with hematoxylin & eosin. For each NHP, one section from the bronchi and one section from each lobe on the right and left lung were selected for scoring, for a total of six sections. Pathologic findings on each slide were scored on a scale of 0 – 2 for intraalveolar edema, the amount of BALT (bronchiolar-associated lymphoid tissue), and the presence of Interstitial inflammation; and 0 – 3 for perivascular inflammatory infiltrates (cuffing), and intraalveolar inflammation. (See Table S1.) Scores for the six slides from each macaque were tabulated, and a cumulative average score was calculated for each NHP using a total of 30 scores as replicates.

### SARS-CoV-2 Viral Genomic and subgenomic RNA quantitative RT-PCR

The presence of viral RNA and viral replication in the BAL, NS, and OS after SARS-CoV-2 P.1 strain challenge was determined by quantitative RT-PCR. RNA was isolated from 200 μL sample using the QIAamp MinElute Virus spin kit (Qiagen, Frederick, MD). For the qRT-PCR assay, viral RNA was first isolated from BAL, NS and OS using the Qiagen MinElute virus spin kit. To generate a control for the amplification reaction, RNA was isolated from the applicable virus stock using the same procedure. The number of copies for the control were calculated using known RNA weights per mol. A master mix was prepared containing Taq-polymerase, obtained from the TaqMan RT-PCR kit (Bioline cat# BIO-78005), RT, RNAse inhibitor, a primer pair at 2 μM concentration (2019-nCoV_N1-F: 5’-GAC CCC AAA ATC AGC GAA AT-3’ and 2019-nCoV_N1-R: 5’-TCT GGT TAC TGC CAG TTG AAT CTG-3’) and probe (2019-nCoV_N1-P: 5’-FAM-ACC CCG CAT TAC GTT TGG TGG ACC-BHQ1-3’) at a concentration of 2 μM. For the reactions, 45 μL of the master mix and 5 μL of the sample RNA were added to the wells of a 96-well plate. All samples were tested in triplicate. Control RNA was prepared to contain 10^6^ to 10^7^ copies per 3 μL. Eight (8) 10-fold serial dilutions of control RNA were prepared and produced a standard curve with a range of 1 to 10^7^ copies/reaction. For amplification, the plate was placed in an Applied Biosystems 7500 Sequence detector and amplified using the following program: 48ºC for 30 minutes, 95ºC for 10 minutes followed by 40 cycles of 95ºC for 15 seconds, and 1 minute at 55ºC. Duplicate samples of each dilution were prepared as described above. If the copy number exceeded the upper detection limit, the sample was diluted as needed. The number of copies of RNA per mL was calculated by interpolation from the standard curve and multiplying by the reciprocal of 0.2 mL extraction volume. This gave a practical range of 50 to 5 × 10^8^ RNA copies per mL for BAL samples and for nasal and oral swabs the viral loads were given per swab.

The RT-PCR assay for the sgRNA utilizes primers and a probe specifically designed to amplify and bind to a region of the N gene mRNA from the Coronavirus, which is not packaged into the virion. The signal was compared to a known standard curve of plasmid containing the sequence of part of the messenger RNA and calculated to give copies per ml. The control DNA was prepared to contain 10^7^ copies. Seven 10-fold serial dilutions of control RNA were prepared using Buffer AVE and generated a standard curve with a range of 1 to 10^6^ copies/reaction. Duplicate samples of each dilution were prepared as described above with the primer pair (SG-N-F: CGATCTCTTGTAGATCTGTTCTC and SG-N-R: GGTGAACCAAGACG CAGTAT) and probe (FAM-TAACCAGAATGGAGAACGCAGTGGG -BHQ). If the copy number exceeded the upper detection limit, the sample was diluted as needed. For amplification, the plate was placed in an Applied Biosystems 7500 Sequence detector and amplified using the following program: 48°C for 30 minutes, 95°C for 10 minutes followed by 40 cycles of 95°C for 15 seconds, and 1 minute at 55°C. The number of copies of RNA per ml was calculated by interpolation from the standard curve and multiplying by the reciprocal of 0.2 ml extraction volume. This gave a practical range of 50 to 5 × 10^7^ RNA copies per mL for harvested samples.

### Statistical analysis

Statistically significant differences between the geometric mean of IgG concentrations or MFI in groups given different vaccine formulations at each timepoint were determined using a two-way ANOVA followed by Dunnett’s Multiple Comparison. Comparisons of PRNT_50_ values between vaccine formulations was done using a Kruskal-Wallis Test followed by a Dunn’s multiple comparisons test. Correlations between IgG or PRNT titers to viral load in the BAL and NS was examined using the non-parametric Spearman’s correlation test. Differences between histopathological scores in different treatment groups was calculated using a one-way ANOVA followed by Dunnett’s multiple comparison test with the cumulative scores of all slides per macaque as replicates. All statistical analysis was completed using Graphpad Prism 9 software (San Diego, CA).

## Supplementary Materials

Fig. S1. Serum Wuhan-Hu-1 RBD-Specific IgG Kinetics

Fig S2. Pre-and Post-Challenge rVSV-SARS-CoV2-S Neutralizing Antibody Response

Fig S3. Individual TCID_50_ Viral load from the BAL and NS

Fig S4. Individual Genomic Viral RNA Load

Fig S5. Individual Subgenomic N transcript RNA Load Table

S1 Detailed Histopathology Scoring System

## Acknowledgements

We would like to thank Takaji Wakita, from the National Institute of Infectious Diseases, Japan, for the P.1 viral challenge strain, and Shelby O’Connor and John Baczenas from the University of Wisconsin-Madison, USA, for deep sequencing our inhouse viral stock. We also would like to thank Adrian Creanga from the Vaccine Research Center-NIAID, USA for the Vero TMPRSS2 cell line used in our TCID_50_ and PRNT_50_ assays, as well as Andrea Marzi, Laboratory of Virology, NIAID-NIH, USA, for the rVSV-SARS-CoV 2 virus stock used in our surrogate PRNT_50_ assay. Furthermore, we would like to thank Melissa Hamilton, the study coordinator at Bioqual Inc., USA; Mapp Biopharmaceutical Inc, for the CR3022 mAb used for protein purification; BTG International Limited for the CoVaccine HT™ adjuvant, and Miyoko Bellinger and Kristen Ewall from the JABSOM Histocore for preparing lung tissue sections for histopathological analysis.

## Funding

This work was supported by:

National Institute of Allergy and Infectious Diseases Grant R01AI132323 (ATL)

National Institute of Minority Health and Health Disparities Grant U54MD007601 (JRH)

National Institute of General Medical Sciences Grant P30GM114737 (RY)

Institutional funds from University of Hawai‘i at Mānoa, BIOQUAL Inc. and Soligenix Inc.

## Author Contribution

Conceptualization: AT, MML, HA, OD, ATL

Methodology: AT, TSW, KT, LP, HA, CYL, ATL

Investigation: AT, TSW, KT, LP, JG, ND, AC, BN, ZF, AVR, JYO

Formal Analysis: AT, MML, KT, ATL

Visualization: AT, KT

Funding acquisition: OD, ATL

Project administration: HA, OD,

ATL Supervision: HA, OD, ATL

Writing – original draft: AT, KT, ATL

Writing – review & editing: AT, TSW, MML, KT, HA, OD, ATL

## Competing Interests

ATL and OD are named inventors on a patent application covering a recombinant subunit vaccine for SARS-CoV-2. LP, JG, ND, AC, BN, ZF, AVR, JYO and HA are current employees of BIOQUAL, Inc. OD is a current employee of Soligenix Inc. All other authors declare no competing interests.

## Data and material availability

All data are available in the main text or the supplementary materials

