## Supplemental Figures and Table for "Protein Vaccine Induces a Durable, More Broadly Neutralizing Antibody Response in Macaques than Natural Infection with SARS-CoV-2 P.1"

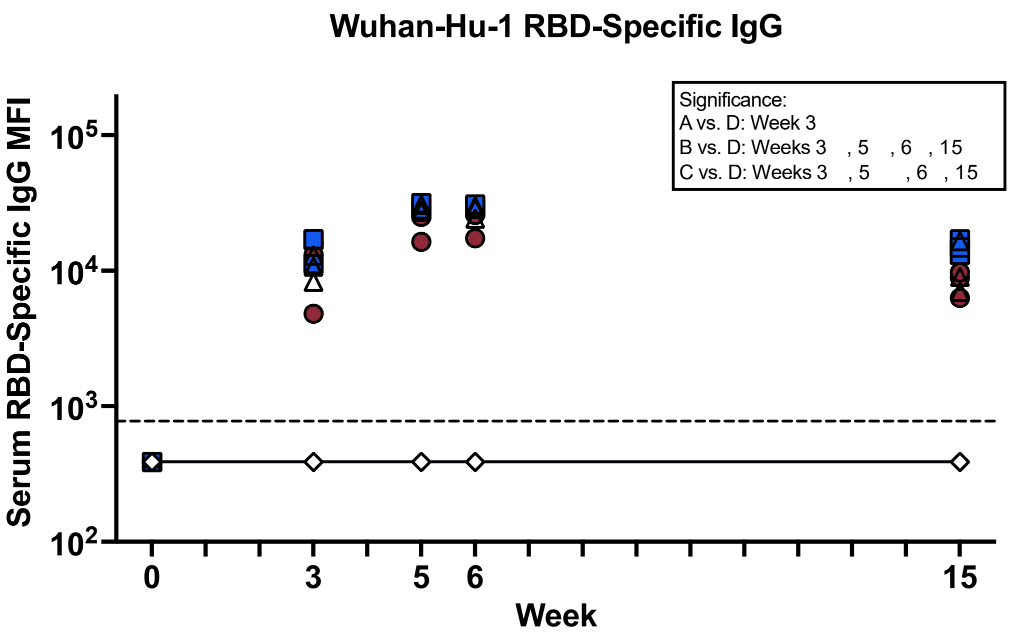


**Figure S1. Serum Wuhan Hu1-Specific IgG Kinetics** Serum S-specific IgG kinetics measured using a MIA. Titers were expressed as MFI. Dashed line indicates the limit of detection (LOD, =733 MFI). Individual values falling below the LOD were set to ½ LOD. Significance was calculated using a one-way ANOVA followed by a Dunnett's Multiple Comparison to Group D at each time point. (*p≤0.05, **p≤0.01)

808

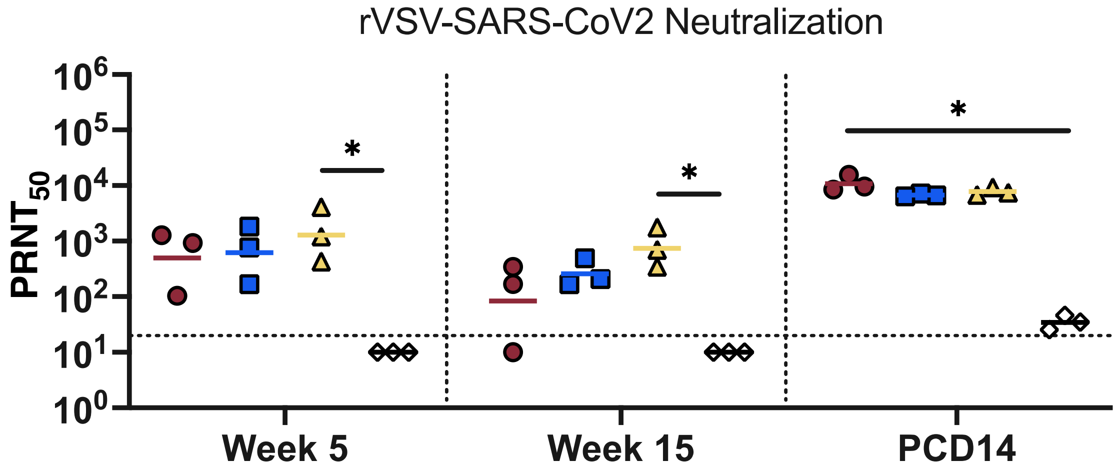


**Figure S2. Pre-and Post-Challenge rVSV-SARS-CoV2-S Neutralizing Antibody Response**
Neutralizing antibodies collected at week 5, week 15 and at 14 days after P.1 challenge were measured using a rVSV-SARS-CoV-2 S PRNT assay. Curve-fitted $PRNT_{50}$ titers were calculated using a sigmoidal dose response curve. Dashed line indicates the LOD (≤1:20). Individual values falling below the LOD were set to ½ LOD. Significance was calculated using a Kruskal-Wallis Test followed by a Dunn's Multiple Comparison to Group D at each time point. (*p≤0.05).

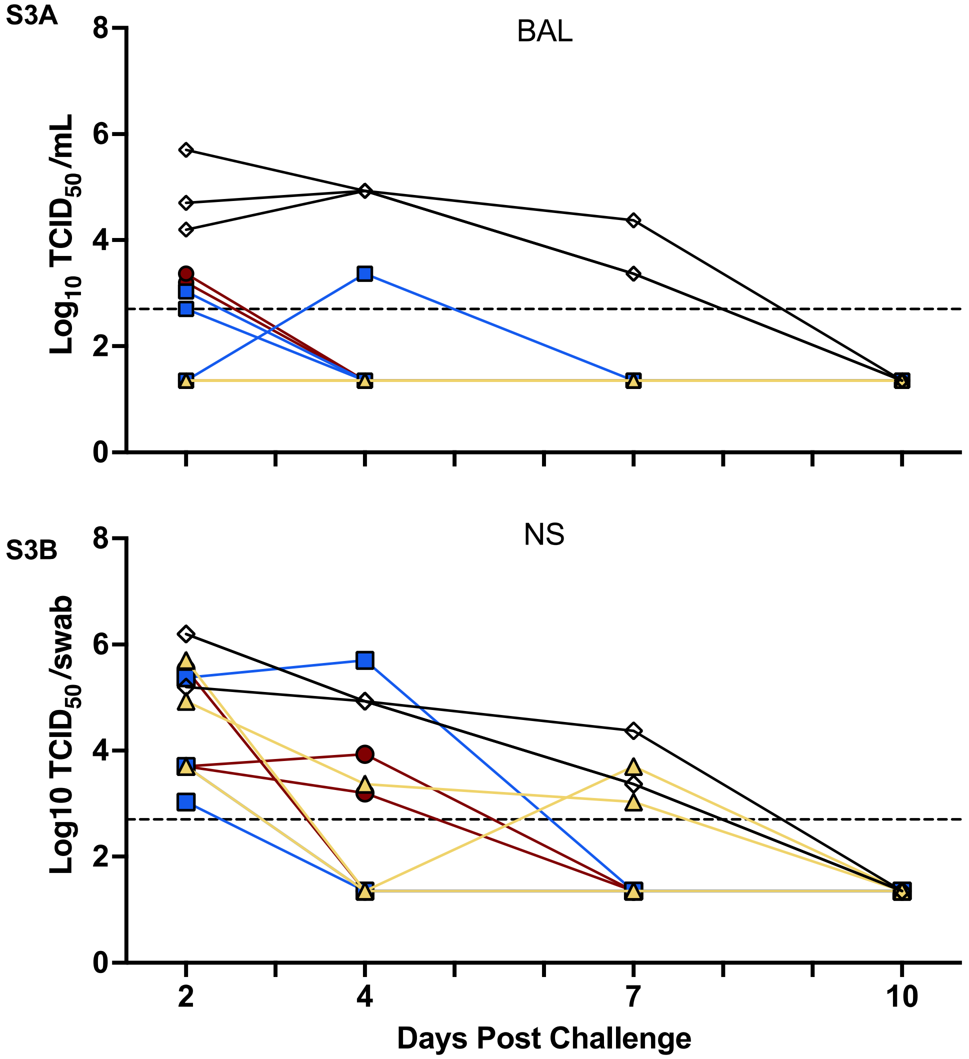


**Figure S3. Individual $TCID_{50}$ Viral Load from the BAL and NS** Serial diluted samples from the (A) lower and (B) upper respiratory tract at each timepoint were incubated with Vero E6 cells and then observed for cytopathology effect (CPE). Endpoint titers were determined at the dilution showing 50% CPE.

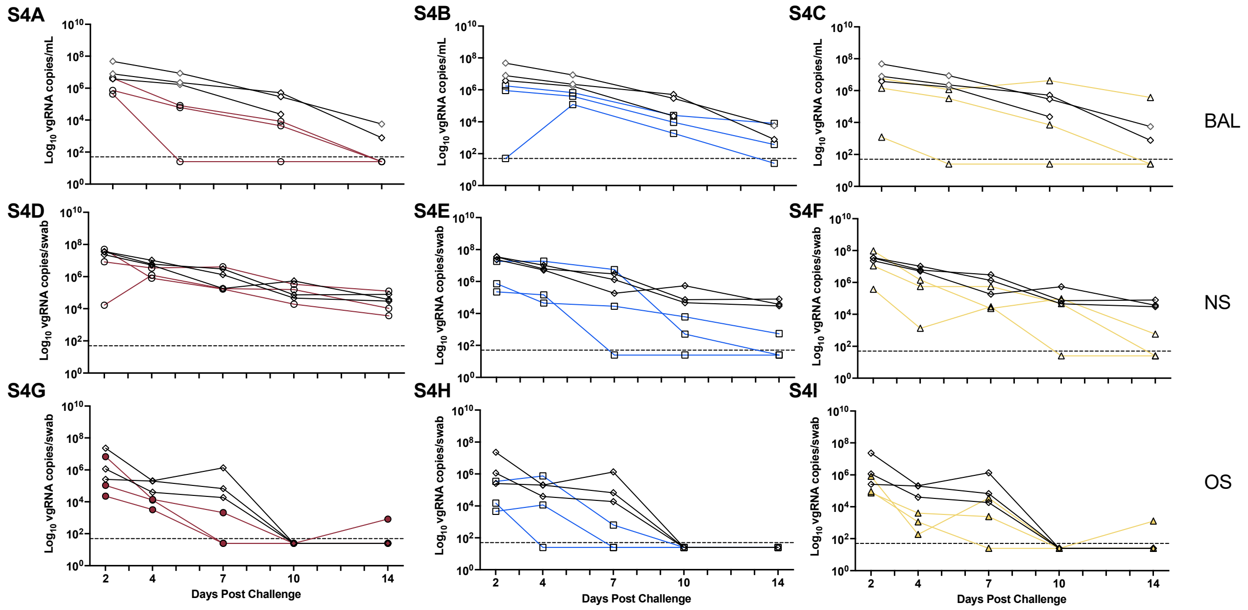


**Figure S4. Individual Genomic Viral RNA Load** Samples were collected from the (A-C) BAL, (D-F) nasal (NS) and (G-I) oral swab (OS) from each macaque at each timepoint and the viral genomic RNA was quantified using qPCR. Individual viral load kinetics from Groups A (A, D, G), B (B, E, H) and C (C, F, I) are plotted together with Group D for comparison. Dashed line indicates the LOD.

811
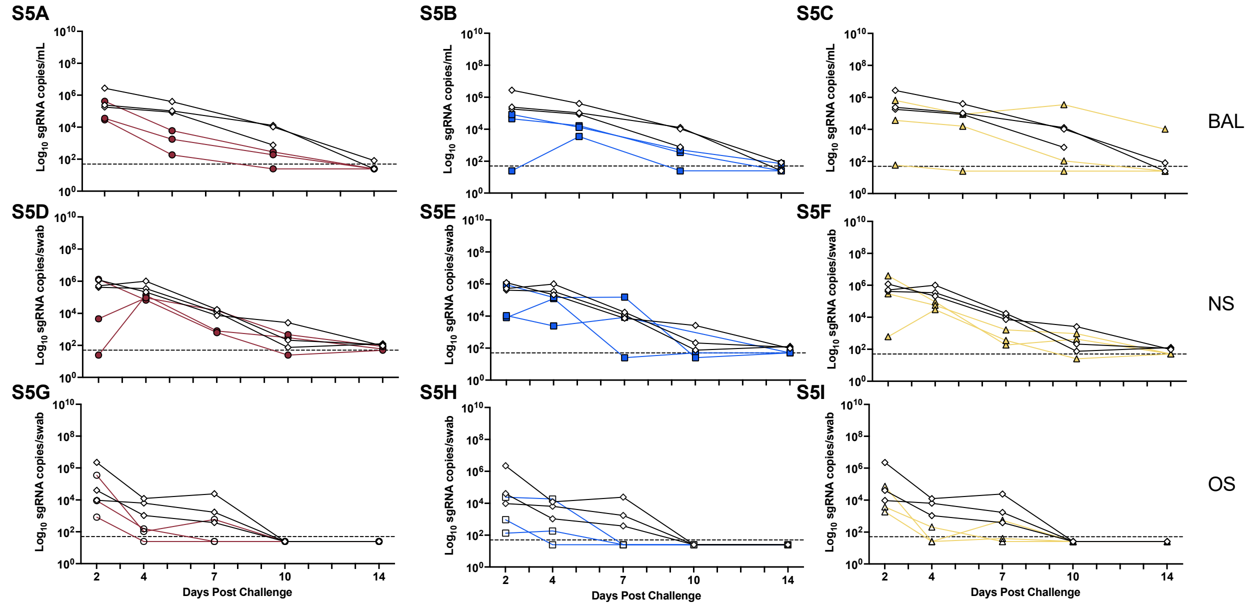


**Figure S5. Individual Subgenomic N transcript Viral RNA Load** Samples were collected from the (A-C) BAL, (D-F) nasal (NS) and (G-I) oral swab (OS) from each macaque at each timepoint and the subgenomic N transcript RNA was quantified using qPCR. Individual viral load kinetics from Groups A (A, D, G), B (B, E, H) and C ( C, F, I) are plotted together with Group D for comparison. Dashed line indicates the LOD.

**Supplemental Table 1: Detailed Histopathology Scoring System**

| | 0 | 1 (minimal) | 2 (mild) | 3 (moderate) |
|---|---|---|---|---|
| **Intraalveolar edema** | Absent | Occasional alveoli, up to 5% of the slide affected | Confluent alveoli; 6-25% of the slide affected | Confluent alveoli; 26-50% of the slide affected |
| **Intraalveolar inflammation** | Alveolar macrophages [a]WNL | Occasional alveoli with increased macrophages | Multiple foci of alveoli with increased or clustered macrophages, and/or multinucleated giant cells | Criteria for 2 with associated acute inflammation (acute pneumonia) |
| **Interstitial inflammation** | Absent | Focal | Multifocal | / |
| **Perivascular inflammatory infiltrates (cuffing)** | None | Occasional, incomplete, or loosely formed cuffs | Numerous cuffs; predominantly incomplete and loosely formed with lesser well-formed complete cuffs | Numerous cuffs; approximately half or more well-formed, and may have a few broad, dense cuffs |
| **Amount of [b]BALT** | [a]WNL | Mildly increased | Moderately increased | / |

[a]WNL: Within Normal Limits. Results are in range with what is considered normal.
[b]BALT: Bronchiolar-associated lymphoid tissue
